# Heptad repeat regions of coiled-coil domains of Mfn1 are crucial for Mfn1 mediated mitochondrial fusion

**DOI:** 10.1101/495721

**Authors:** Sansrity Sinha, Aradhyam Gopala Krishna

## Abstract

Mitofusin mediate fusion of outer mitochondrial membranes (OMM). Recent studies have explained the role of GTPase domain for Mfn1 dimerization and outer mitochondrial membrane (OMM) fusion. Coiled-coiled domains [namely, coiled-coil/Middle domains (CC1MD) and coiled-coil-2 GTPase effector domain (CC2/GED)] form helical bundles that mediate open-to-close conformations of Mfn1 upon GTP binding and have been previously reported to be important for OMM tethering and OMM fusion. To further decipher the significance of helical structure of MD, we functionally characterized the heptad repeat regions of MD. Consistent with previous studies, we show that MD consists of two heptad repeats (HR1, namely HR1a and HR1b) and both of these are crucial for Mfn1 mediated OMM fusion.

**HIGHLIGHTS:**

- Coiled-coil1 (also known as Middle domain, MD) contains two heptad repeat regions.
- Heptad repeats of MD (namely HR1a and HR1b) are crucial for fusogenic property of Mfn1
- Mutations disrupting of helical structure of HR1b lead to loss of fusogenic activity of Mfn1

## 1. INTRODUCTION

Proteins regulating organellar membrane remodeling and dynamics belong to dynamin superfamily[1–4]. In general, all dynamin superfamily proteins (DSPs) have a conserved GTPase domain and two conserved coiled-coil domains which mediate open-close conformations upon ligand binding and hydrolysis and thereby the function[1–6]. Mitochondrial dynamics is mediated by a subset of DSPs, known as dynamin-related proteins (DRPs) that regulate mitochondrial size and number within the cell. Mitochondrial fission is mediated by Dynamin-related protein 1 (Drp1), which is a cytoplasmic protein and is recruited to outer mitochondrial membrane (OMM) by means of adapter proteins for mediating mitochondrial division/fission. Mitochondrial fusion is mediated by DRPs that localize on mitochondrial membranes. Mitofusins (Mfn1 and Mfn2, also known as fuzzy onions, Fzo) mediate OMM fusion while optic atrophy-1 (Opa1) mediates inner mitochondrial membrane (IMM) fusion[1–16].

Recent studies have explained the role of GTPase domain for Mfn1 dimerization and outer mitochondrial membrane (OMM) fusion. Coiled-coiled domains [namely, coiled-coil/Middle domains (CC1MD) and coiled-coil-2 GTPase effector domain (CC2/GED)] form helical bundles that mediate open-to-close conformations of Mfn1 upon GTP binding and have been previously reported to be important for OMM tethering and OMM fusion[10, 15–17]. Here we characterize MD/CC1 of Mfn1 to further decipher the significance of its helical structure in Mfn1 mediated OMM fusion.

## 2. METHODS

### 2.1 Data retrieval and determination of domain architecture

NCBI was used as a source of peptide sequences used for the study. Homolog mining was done using NCBI BLAST tool (BLASTp, BLASTn, and tBLASTn strategy) (http://blast.ncbi.nlm.nih.gov/Blast.cgi) to identify all annotated and un-annotated sequences used in the study. Only full-length sequences were taken up for further studies.. The homologs were also confirmed for the domain architecture search using Pfam (http://pfam.xfam.org/search) and NCBI CDD (http://www.ncbi.nlm.nih.gov/Structure/cdd/wrpsb.cgi). Also, COILS (http://embnet.vital-it.ch/software/COILS_form.html) and TargetP (http://www.cbs.dtu.dk/services/TargetP/) [18, 19] available at Expasy were used to determine coiled-coil regions and the subcellular localization of the protein. The sequences used in this study are enlisted in Table.S1.

### 2.2 Multiple sequence alignment and helical wheel plot generation

Multiple sequence alignment (MSA) was performed for identified Mfn1 and Mfn2 orthologs in euteleostomes using MAFFT v.7.25 (G-INS-1 strategy, BLOSUM62 matrix including gappy regions, 10,000 maxiterate, with threshold score =39, E value = 8.4e-11 and Gap opening penalty of 1.53 with an offset of 0.5) using a total of 32 sequences (Supplementary File.S1). Draw coil1.0 was used to draw helical wheel plots of predicted HR1b of human Mfn1 and Mfn2 sequences (https://grigoryanlab.org/drawcoil/).

### 2.3 Structural modeling

The human Mfn1 protein was modeled using i-TASSER server (http://zhanglab.ccmb.med.umich.edu/I-TASSER/) [20–22]. The best fit model for both proteins was selected based on C-score and based on further examination of the stereo chemical parameters by SAVES server (SAVES v5.0) (https://www.acronymfinder.com/Structure-Analysis-and-Verification-Server-(University-of-California%2c-Los-Angeles)-(SAVΈS).html). FATCAT was used for superposition of modeled structure of human Mfn1 with the crystal structure of Mfn1 (PDB: 5GOF).

### 2.4 DNA manipulations and cloning

For this purpose, we cloned Mfn1 in pDsRed2n1 between BamHI and XhoI to generate WtMfn1-pDsRed2n1 construct. Using Stratagene QuickChange site directed mutagenesis kit we generated L380P, L380/390P and ΔCC1-Mfn1-dsRed2n1 constructs of Mfn1. These constructs were used for functional assays in HEK293T cells.

### 2.5 Cell culture, maintenance and transfections

HEK293T and MCF7 cells were obtained from NCCS, India. Cells were maintained in DMEM supplemented with 10% FBS in 5% CO_2_. 3 x 10^5^ cells were seeded 24hrs before transfection and were grown upto 70% confluency. Transfection was done using lipofectamine 3000 as per standard manufacturer’s instructions. 24hrs prior to transfection, 1 x 10^5^ cells were seeded per 35mm dish. Transfection was done using Lipofectamine 3000 using standard manufacturer’s instructions. 1μg of total DNA was used for transfection per 35mm dish. Images were taken using Olympus IX51 microscope 48hrs after transfection. Post-imaging cells were used for western blot analyses.

### 2.6 Lysate preparation and western blot analyses

Cells were lysed in Radioimmunoassay precipitation buffer (RIPA) with protease inhibitor cocktail (Roche life science) to prepare lysates. The lysates were used to probe expression profile of proteins by western blot. 1:1000 dilution of the following primary antibodies in 3% BSA was used: Mfn1 (Cell signalling technology) and Actin (Santa Cruz Biotechnology). Following secondary antibodies tagged with horse radish peroxidase were used at a dilution of 1:5000 – anti-rabbit (Santa Cruz Biotechnology), anti-mouse (Santa Cruz Biotechnology).

## 3 RESULTS AND DISCUSSION

### 3.1 Deletion of MD/CC1 leads to production of degradation of Mfn1 due to improper folding of protein

In order to decipher the functional relevance of MD/CC1, we deleted this region in Mfn1-pDsRed2n1 plasmid (Fig.1c). Upon transfection of WtMfn1 and of ΔCC1-Mfn1, we found that WtMfn1 localizes in the mitochondria while the deletion mutant of coiled coil 1 domain failed to express (Fig1.d). Thus, it is plausible to assume that MD/CC1 is crucial to mediate Mfn1 mediated OMM fusion, as shown in previous studies [10, 15–17, 23]. It is plausible to assume that the deletion of MD/CC1 either hampers the structure of Mfn1 (and henceforth its function) or interferes with the folding of Mfn1thereby inhibiting the fusogenic activity of Mfn1.

**Fig.1:**
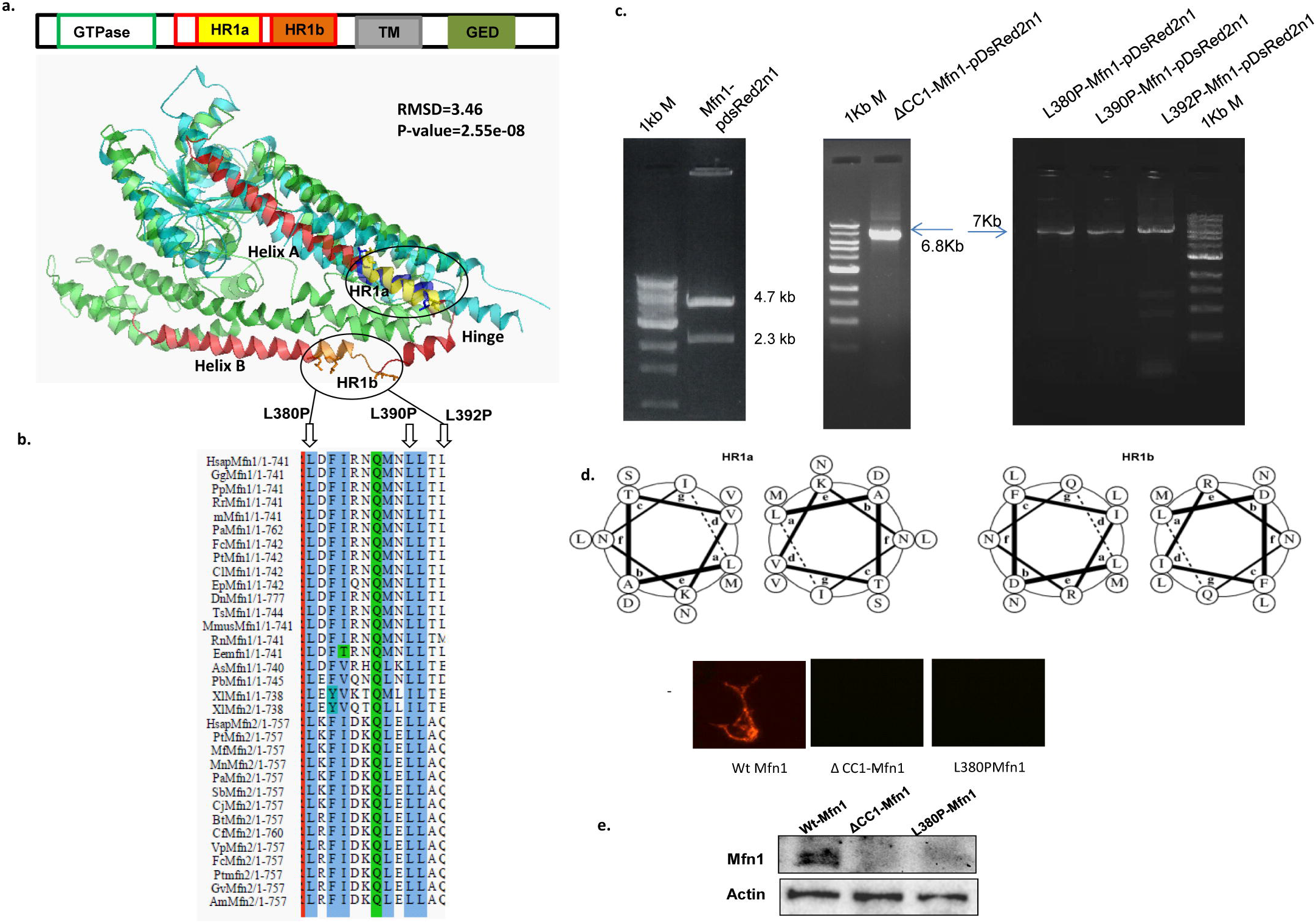
Identification and characterization of heptad repeat regions in MD/CC1: a. Schematic representation of domains of Mfn1. Superposition of Mfn1 structures (5GOF and modeled structure) using FATCAT server. The MD/CC1 is shown with red color. The heptad repeats (HR1a and HR1b) are shown with yellow and orange colors, respectively. The HR1a in the crystal structure is shown with dark blue color. The conserved leucines marking the terminals of both heptad repeats are shown by sticks. b. The HR1b alignment is shown for a Mfn1 and Mfn2 sequences showing the conserved leucines (Leu380, L390 and Leu392). The corresponding helical wheel plots of HR1a and HR1b are shown in (d). c. Gel image showing double digestion of Mfn1 cloned in pDsredn1 with BamHI and XhoI showing release of 2.3Kb Mfn1 fragment. Gel image showing the generation of Leu380Pro, Leu390Pro and Leu392Pro in WtMfn1 using site directed mutagenesis. e. Fluorescence microscopy images mitochondrial localization of Mfn, ΔCC1-Mfn1 and Leu380Pro-Mfn1 (left panel) and EGFP (right panel). The corresponding western blots showing the band for Mfn1 and is shown. Actin was used as loading control.

### 3.2 The conserved Heptad repeats of MD/CC1 lie on either helices of MD in Mfn1 and Mfn2 sequences

The results from MSA suggest that the presence of two heptad repeats in the MDCC1 region of Mfn1/2 sequences. Both HR1a (Leu348-Leu360) and HR1b (Leu380-Leu392) are conserved across the euteleostome Mfn1/2 sequences (Fig.1a). The helical wheel plots suggest that both HR1a and HR1b regions in Mfn1 and Mfn2 forms an amphipathic helices. Since the reported crystal structures (PDB ID: 5go4, Apo form) lack the HR1b, the TM (substituted by artificial Ser-Ala linker) and the interdomain regions, the structure of human Mfn1 and Mfn2 were modeled and were evaluated for stereo chemical parameters using SAVES server (Supplementary table S1) [10, 17]. The modeled structure closely resembles the reported structures (5go4 vs. model4 HsMfn1, P-value= 2.55e-08, RMSD=3.46, raw score=678.96, without twists using rigid model, FATCAT server). To identify the structural basis for observed results and gain structural insights for the topological location of this region in the structure of Mfn1, and this region was mapped onto the modeled structure. The sequence represented a helical region in the Helix B of MD/CC1 (Fig.1b).

### 3.3 The conserved HR1b is crucial for Mfn1 mediated OMM fusion

We rationalized that MD/CC1 may be important for structural integrity of Mfn1 and therefore its function, so we generated mutants that disrupt the helical structure of MD/CC1. We observed that apart from high conservation of initial (Leu380) and terminal (Leu392) leucines, all Mfn1 sequences had an additional conserved leucine (Leu390) that was completely absent in Mfn2 sequences. Consistent with characterized mutations suggested for HR2 of GED, Leu to Pro mutants of HR1b were generated. Thus, we generated point mutants of Leu380P, Leu390P and Leu392P (Fig.1c). Upon co-transfection of WtMfn1 and all three Leu-Pro mutants with empty pEGFPn1, similar to ΔCC1-Mfn1 the mutants did not express protein (Fig.1e). Hence, we propose that the structural integrity of HR1b is crucial for Mfn1 mediated OMM fusion.

Interestingly, HR1b lies adjacent to previously characterized HR1 region. This region (Thr393-Ser10) was also shown to bind to membrane and crucial for Mfn1 mediated OMM fusion [23]. Furthermore, collating our results with previous studies, we hypothesize that this region might play a role in mediating open-to-close conformation of Mfn1 following GTP binding, serving as internal conformation-switch in Mfn1.

## 4 CONCLUDING REMARKS

In summary, consistent with previous studies, we show that MD consists of two heptad repeats (HR1, namely Hr1a and HR1b) and both of these are crucial for Mfn1 mediated OMM fusion. Our results demonstrate that MD crucial for Mfn1 mediated mitochondrial fusion. Also, consistent with reported HR2 mutations that disrupt its helical structure, mutations disrupting the helical structure of HR1b lead to loss of fusogenic property of Mfn1[10, 15–17, 23]. Hence, we hypothesize that the heptad repeats of MD and GED domains are important for Mfn1 mediated OMM fusion. Since HR1a has been shown previously to be crucial for Mfn1 mediated OMM fusion, collating our results with previous studies we propose that the interaction between HR1a and HR1b is probably necessary for maintaining mitofusin in inactive or GDP bound, or closed state. These interactions are broken upon GTP binding, (converting mitofusin from closed to open state conformation) thus providing transition between the open and closed state confirmations of mitofusin that enables mitochondrial membrane tethering and fusion. Together we demonstrate that coiled-coil domains are not just importance for maintaining helical structure of mitofusin but also act as a conformational switch within the protein thus enabling its function.

## 5. ACKNOWLEDGMENTS

The authors acknowledge financial support from IIT Madras and infrastructure support from the Bioinformatics Infrastructure Facility and Central Equipment facility, IIT Madras.

## ABBREVIATIONS

DSPs: (Dynamin Superfamily proteins)
DRPs: (Dynamin-related proteins)
HR1: (Heptad repeat 1)

